# Segmentation of Glomeruli Within Trichrome Images Using Deep Learning

**DOI:** 10.1101/345579

**Authors:** Shruti Kannan, Laura A. Morgan, Benjamin Liang, McKenzie G. Cheung, Christopher Q. Lin, Dan Mun, Ralph G. Nader, Mostafa E. Belghasem, Joel M. Henderson, Jean M. Francis, Vipul C. Chitalia, Vijaya B. Kolachalama

**Affiliations:** Section of Computational Biomedicine, Department of Medicine, Boston University School of Medicine, Boston, MA; College of Engineering, Boston University, Boston, MA; College of Health & Rehabilitation Sciences, Sargent College, Boston University, Boston, MA; Renal Section, Department of Medicine, Boston University School of Medicine, Boston, MA; Department of Pathology and Laboratory Medicine, Boston University School of Medicine, Boston, MA; Whitaker Cardiovascular Institute, Boston University School of Medicine, Boston, MA; Veterans Administration Boston Healthcare System, Boston, MA; Hariri Institute for Computing and Computational Science & Engineering, Boston University, Boston, MA

**Author notes:** Corresponding author: Vijaya B. Kolachalama, PhD, Boston University School of Medicine, 72 E. Concord Street, Evans 636, Boston, MA, USA – 02118.

**Keywords:** Kidney biopsy, Glomerulus, Digital pathology, Image segmentation, Deep learning, Computational pathology, Trichrome stain

## Abstract

**Introduction:** The number of glomeruli and glomerulosclerosis evaluated on kidney biopsy slides constitute as standard components of a renal pathology report. Prevailing methods for glomerular assessment remain manual, labor intensive and non-standardized. We developed a deep learning framework to accurately identify and segment glomeruli from digitized images of human kidney biopsies.

**Methods:** Trichrome-stained images (n=275) from renal biopsies of 171 chronic kidney disease patients treated at the Boston Medical Center from 2009-12 were analyzed. A sliding window operation was defined to crop each original image to smaller images. Each cropped image was then evaluated by three experts into three categories: (a) No glomerulus, (b) Normal or partially sclerosed glomerulus and (c) Globally sclerosed glomerulus. This led to identification of 751 unique images representing non-glomerular regions, 611 images with either normal or partially sclerosed (NPS) glomeruli and 134 images with globally sclerosed (GS) glomeruli. A convolutional neural network (CNN) was trained with cropped images as inputs and corresponding labels as output. Using this model, an image processing routine was developed to scan the test data images to segment the GS glomeruli.

**Results:** The CNN model was able to accurately discriminate non-glomerular images from NPS and GS images (Performance on test data - Accuracy: 92.67±2.02% and Kappa: 0.8681±0.0392). The segmentation model that was based on the CNN multi-label classifier accurately marked the GS glomeruli on the test data (Matthews correlation coefficient = 0.628).

**Conclusion:** This work demonstrates the power of deep learning for assessing complex histologic structures from digitized human kidney biopsies.

## INTRODUCTION

Glomerular damage is a common manifestation in a spectrum of renal diseases that lead to chronic kidney disease (CKD) and end stage renal disease (ESRD) [1]. Morphological and ultrastructural alterations within the glomeruli provide invaluable information on the mechanisms of renal impairment and facilitate accurate clinical diagnosis [1–3]. Assessment of this highly relevant structure is therefore integral to histopathological analysis of kidney biopsies. A fundamental morphologic parameter in nephropathology is the quantification of normal and abnormal glomeruli in the light microscopic material. For example, number of glomeruli is required for assessment of tissue sufficiency in kidney transplant pathology. Also, histological analysis of glomerular diseases involves careful examination of the entire kidney biopsy slide, and this includes in part, identification of all the glomeruli, assessment of the state of each glomerulus, and integration of this data with other parameters to pinpoint the diagnosis of the glomerular disease [4–7]. While this multistep process of counting and assessing all the glomeruli can be handled efficiently at large medical centers under the supervision of an in-house nephropathologist, this expertise is not available at all locations across the globe. In addition, even for the institutions that have the expert nephropathologist, we need approaches that can automatically perform some of these tasks in order to assist clinical practice to maximize their efficiency.

Machine learning (ML), a powerful technique that is increasingly being utilized in medicine, has the ability to perform these tasks in an efficient fashion. ML approaches give computers the ability to integrate discrete datasets in an agnostic manner to detect previously indecipherable patterns and generate a disease-specific fingerprint. ML can leverage many images as inputs and correlate patterns and features with clinical outcomes. Building on the advances of ML, scientists recently have developed so-called “deep learning” frameworks such as convolutional neural networks (CNN) for object recognition and classification [8]. We and others have leveraged these unbiased, self-learning approaches for key renal pathological features such as fibrosis [9]. We now demonstrate a framework to automatically identify and segment normal or partially sclerosed (NPS) glomeruli as well as the glomeruli with global sclerosis (GS) present within digitized images of human kidney biopsies.

## MATERIALS AND METHODS

### Data collection

Anonymized kidney biopsies were obtained and digitized after approval by the Boston University Medical Campus’ institutional review board under waiver of consent (H-26367). Kidney biopsy procedures were performed on selected patients treated at BMC between January 2009 and December 2012 (Table 1). In total, 171 kidney biopsy slides were available for subsequent imaging. These biopsy samples were obtained from adult patients who had a native or an allograft biopsy, independent of the indication for the biopsy procedure [9]. The criterion for inclusion was the availability of pathology slides.

**Table 1.** Patient and digitized kidney biopsy characteristics used for this study.

| **<u>Characteristic</u>** | **<u>Value</u>** | **<u>Units</u>** |
| --- | --- | --- |
| Number of patients | 171 |  |
| Age, median (range) | 52 (19-86) | years |
| Male | 59.6 | % |
| Patients per race/ethnicity (White, Black, Hispanic, Other) | 46, 79, 24, 22 |  |
| BMI, median (range) | 28.94 (15-56.2) | kg/m<sup>2</sup> |
| Creatinine, median (range) | 2.31 (0.54-13.29) | mg/dl |
| eGFR, median (range) | 30 (5-163) | ml/min per 1.73 m<sup>2</sup> |
| Proteinuria, median (range) | 1.79 (0.03-20.5) | g/g |
| Number of unique images | 275 |  |
| Total number of glomeruli | 745 |  |
| Number of normal or partially sclerosed glomeruli | 611 |  |
| Number of globally sclerosed glomeruli | 134 |  |

### Imaging

Biopsy samples were obtained in the form of individual trichrome-stained slides prepared from formalin-fixed, paraffin-embedded core-needle biopsy tissue. A selected core visible on each slide was imaged at 40× magnification (indicating a 4x objective and a 10× eyepiece) using a Nikon Eclipse TE-2000 microscope (Melville, NY; http://www.bumc.bu.edu/busm/research/cores/). Images were generated with a special consideration to cover the entirety of the biopsy sample which resulted in multiple 40× images per patient. All the images were manually focused using the NIS-Elements AR software (Nikon, Tokyo, Japan) that was installed on the computer connected to the microscope. A total of 275 unique images (∼2 40× images per patient) were used and the average size of each image was about 2560×1920×3 pixels, corresponding to a field of 2.176×1.632 mm^2^, which resulted in a length scale of 0.85 µm/pixel. These images were then converted to 8-bit red−green−blue (RGB) color images in TIFF format.

### Glomerular dataset generation

Each of the digitized images represent a large portion of the digitized biopsy (Figure S1), and the information contained within them had to be filtered in order to train a glomerular image classifier. Therefore, we created a dataset that was more amenable for CNN model training using the sliding window operator. This operation allowed us to systematically crop the original images (n=275) into smaller ones of size 300×300×3 pixels. As a result, we obtained 745 images with glomeruli and 751 images with non-glomerular regions, each of size 300×300×3 pixels, which were then used for model training. See **Supplemental material** for more details.

### Model training for glomerular classification

We used Google’s Inception v3 CNN architecture, which was pre-trained on millions of images with 1000 object classes [10], incorporated minor changes to fine-tune the framework and trained it to predict the presence or absence of a glomerulus within the cropped trichrome images (Figure 2). Specifically, we removed the final classification layer from the network and retrained it with our dataset using the 3 output labels (No glomerulus, normal or partially sclerosed (NPS) glomerulus, and glomerulus with global sclerosis (GS)). We then performed fine-tuning of the parameters at all layers. This procedure, known as transfer learning, is optimal, given the amount of data available. See **Supplemental material** for more details.

**Figure 1.**
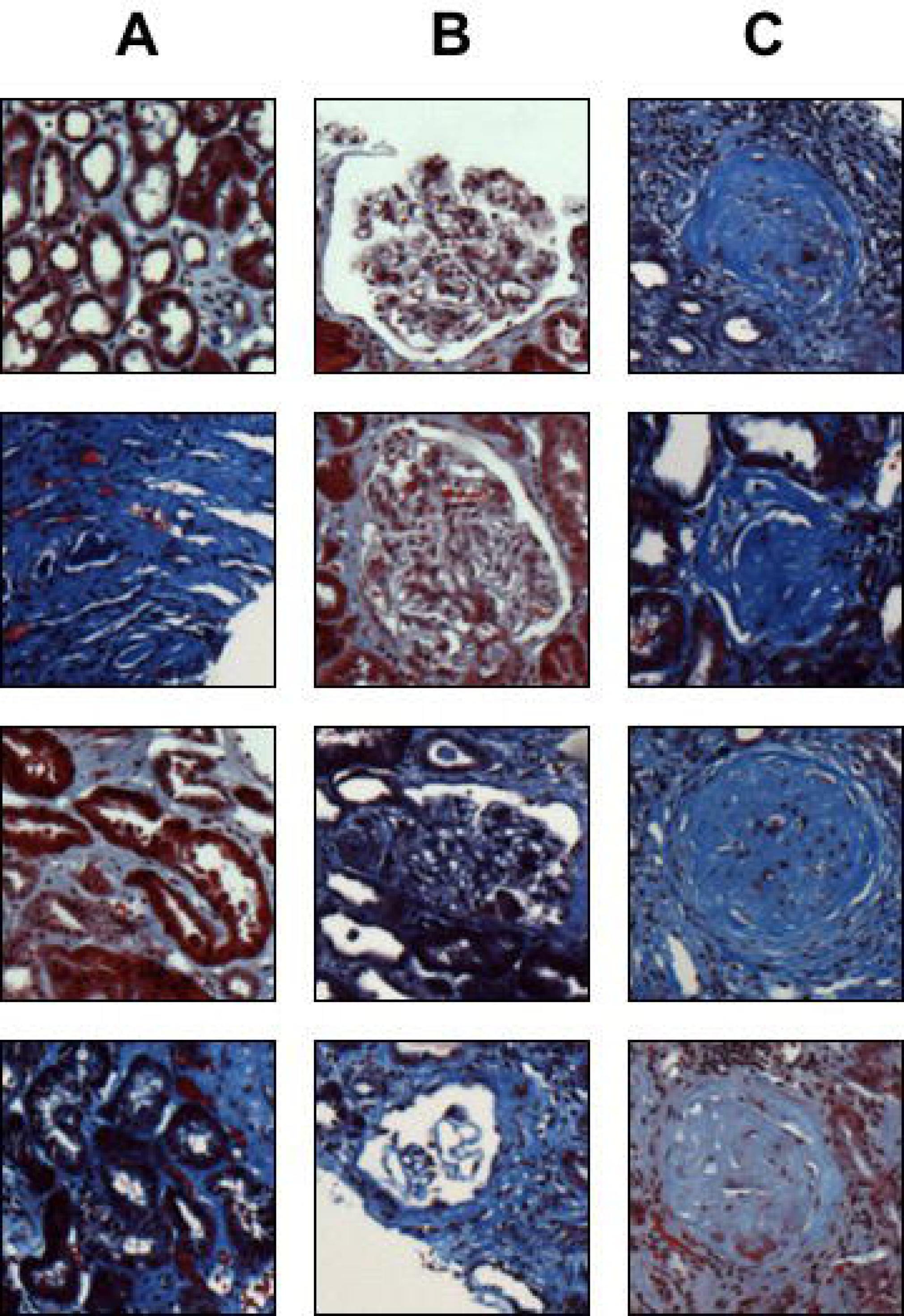
Cropped images. The sliding window operator was used to generate different sets of images to train the CNN model. The first column contains images with non-glomerular tissue, the second column contains images with either a single normal or partially-sclerosed glomerulus, and the third column contains images with a single globally-sclerosed glomerulus. Each cropped image is of size 300×300×3 pixels.

**Figure 2.**
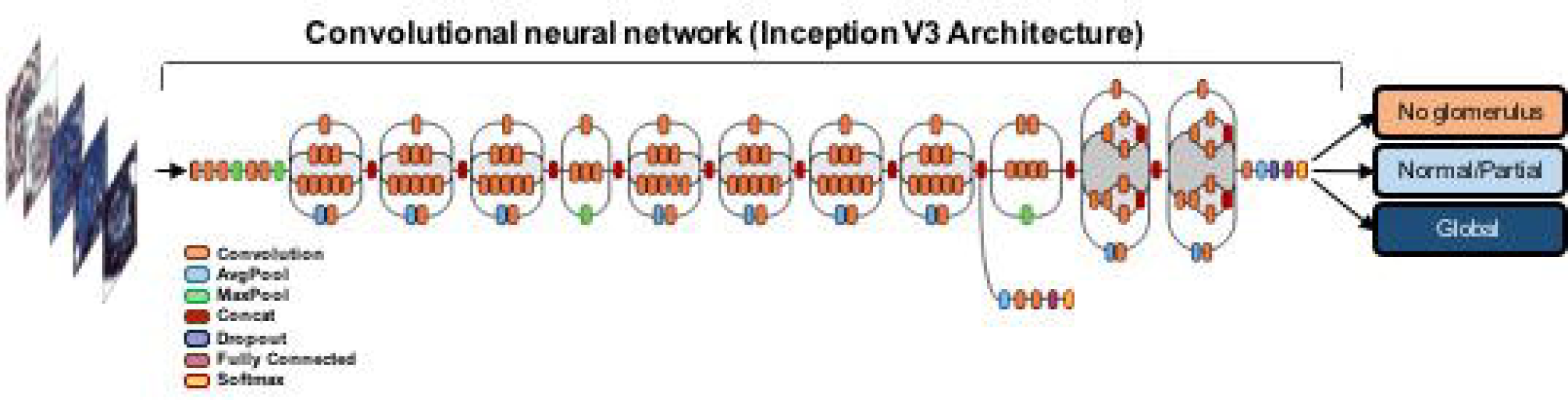
Schematic of the deep neural network. Our classification technique is based on leveraging a pre-trained convolutional neural network (CNN), which was fine-tuned on our dataset (see Methods). The architecture is reprinted with permission (https://research.googleblog.com/2016/03/train-your-own-image-classifier-with.html).

The cropped image dataset was randomly split at the patient level. Specifically, in order to capture intra-and inter-patient variabilities, and to verify whether the CNN model works well on images and image characteristics which it has not been trained on, the patient list was randomly split into 2 parts in a 7:3 ratio (70% training, 30% testing). This resulted in 120 patients in the train set, and 51 patients in the test set. Cropped images belonging to each patient in the list were included in the corresponding dataset (training vs testing). Also, for consistency, we repeated the process of random splitting 4 times. CNN model training and testing were performed on each split, and average performance values were recorded.

### Data augmentation

Some of the glomeruli on the biopsy images were observed on the edges of the tissue sample. When cropping was performed to capture these cases, a portion of the cropped region had only the background pixels. All these images were used as part of the training data, but they were not in sufficient number to be able to generate a model that could accurately identify the glomeruli present on the edges of the biopsy. We therefore augmented the training data by creating copies (n=5) of each image by randomly whitening a small fraction (=0.2) of the total pixels in the images, resulting in 6 total images per original cropped image (Figure 3). See **Supplemental material** for more details.

**Figure 3.**
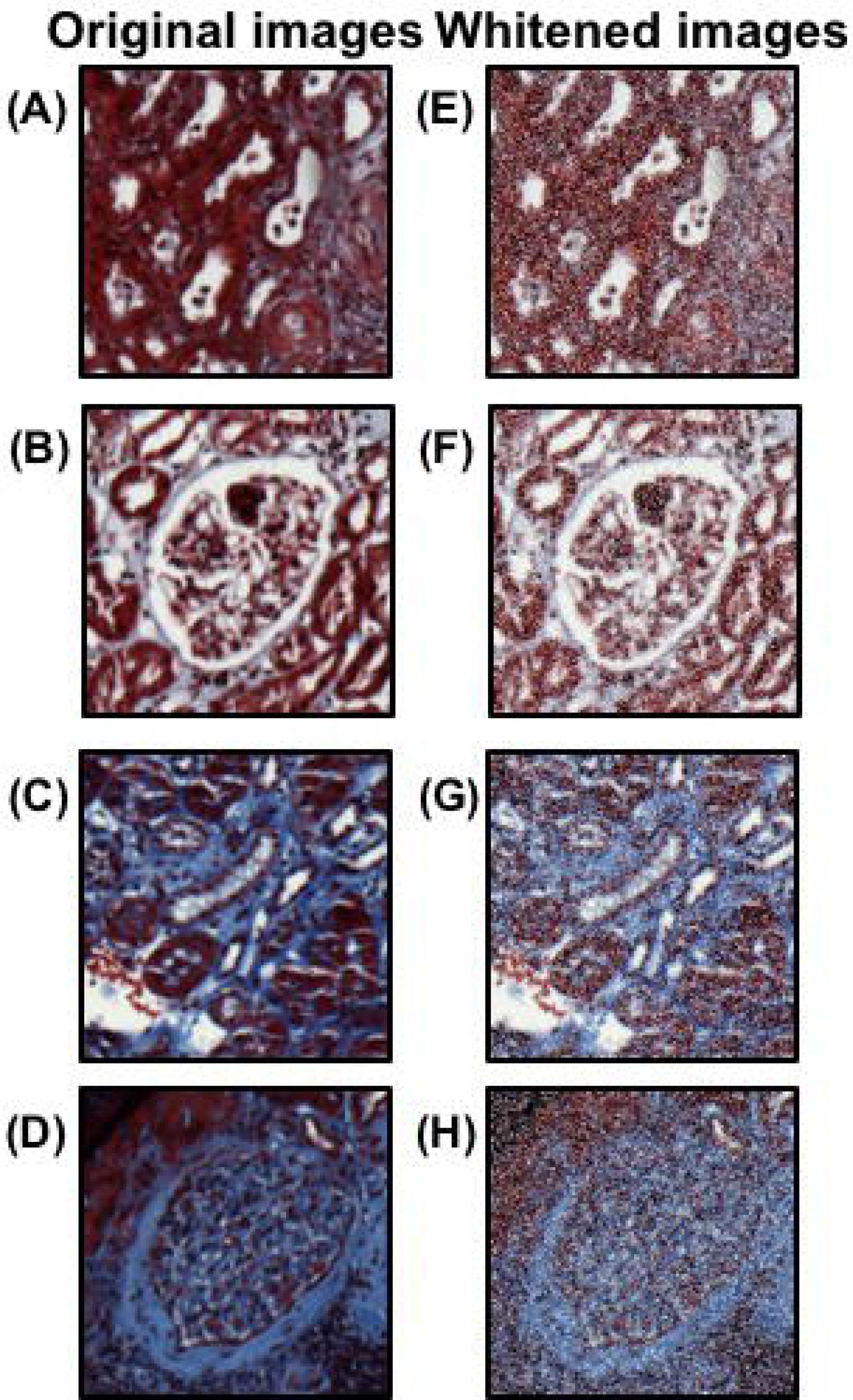
Whitening transformation used for data augmentation. On each cropped image with a single glomerulus (A, B) or a non-glomerular aspect (C & D) of the kidney biopsy, about 20% of the pixels were randomly selected and a whitening transform was applied on them. This process generated images that still contained a major portion of the original content that represented either the glomerular or non-glomerular aspects of the kidney biopsy (E, F, G & H).

### Image segmentation

Using the CNN model with the best test performance, we tested an image processing routine to scan the test images to identify and segment the GS glomeruli (Figure 4). The sliding window operation was used again to scan the entire test image of size ∼2560×1920×3 pixels in increments of 300×300×3 pixels. Each cropped image was then processed through the trained CNN model that predicted if there was a GS glomerulus. An output of ‘0’ indicated that the CNN model determined that no glomerulus was present whereas an output of ‘2’ indicated a GS glomerulus was detected within that cropped image. Note that an output of ‘1’ was reserved for identifying an NPS glomerulus but this result was not used for glomerular segmentation. When a GS glomerulus was detected, the pixel coordinates of the four corners of that image were stored in an array. This process was repeated as the sliding window operation swept from one end of the corner to the other, which resulted in bright patches that corresponded to the areas that were predicted to contain a GS glomerulus. A heatmap was generated using these corners. The brightness of the patch in the heatmap was found to be directly proportional to how confident the model was in terms of detecting the presence of a GS glomerulus in that area. Every non-bright region (i.e., area with pixel intensity close to 0) on the heatmap then represents all the non-glomerulus regions.

**Figure 4.**
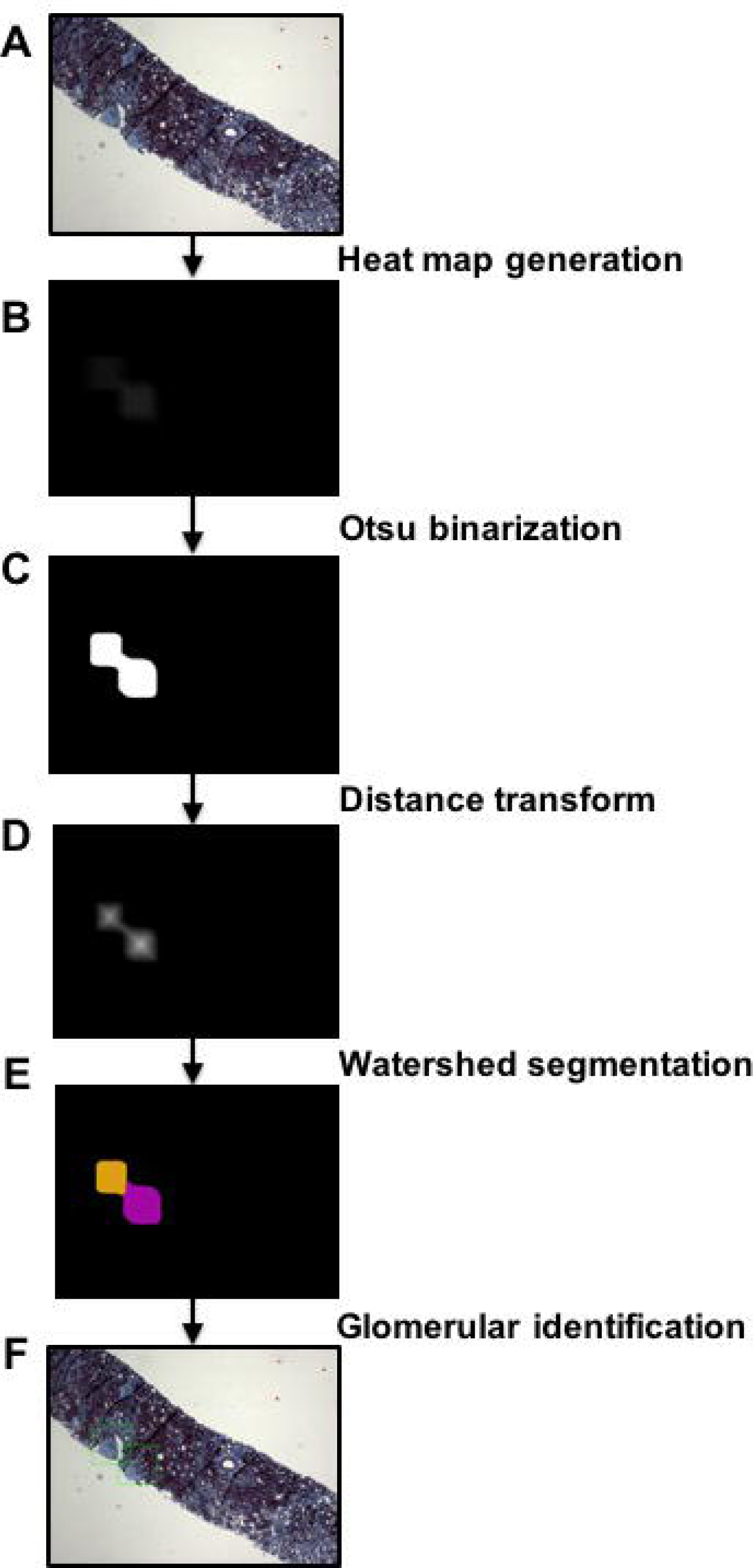
Glomerular segmentation pipeline. The trained CNN model was used in conjunction with the sliding window operator to scan a test image (A) that was not used in model training. (B) A heat map was generated based on how the CNN model detected the presence of GS glomeruli. (C) An Otsu binarization operation was attempted on the heatmap, followed by a distance transform (D) and then watershed segmentation (E), which resulted in segmentation of two distinct GS glomeruli (F).

Generated heatmaps were processed further to segment the identified GS glomeruli using a simple annotation defined as a ‘green box’ surrounding the GS glomerular region. We performed this task by first binarizing the image using the Otsu’s method [11]. Note that the threshold value for binarization was empirically determined (=20), after examining several images. Subsequently, a distance transform was applied on the heatmaps, which simply calculated the distance of each foreground pixel from the nearest background pixel. We then performed watershed segmentation to separate the identified ‘blobs’ in the image. The watershed transformation treats the image it operates upon like a topographic map, with the brightness of each pixel representing its height, and finds the lines that run along the tops of ridges [12]. Finally, a box was automatically placed by the segmentation algorithm to highlight the identified GS glomerulus.

### Performance metrics

Datasets for training and testing were divided such that none of the images belonging to patients in the testing set were available to the model while training. This implies that the training-testing split was done at the patient level as opposed to the image level and the variability in the test images was not part of the images which the model was trained on. This strategy allowed us to systematically evaluate the model performance on completely new patient data. The random splitting of the train and test data was performed 4 different times and model performances on the test data were averaged across all the runs. The CNN model developed to perform multi-label classification was evaluated by computing overall mean accuracy and mean Cohen’s kappa (*k*). The segmentation model performance was evaluated by computing overall accuracy, sensitivity and specificity on the test data. We also computed F_1_-score as a measure of model accuracy that considers both the precision and recall of a test. We also computed Matthews correlation coefficient (MCC), which is a balanced measure of quality for dataset classes of different sizes of a binary classifier.

## RESULTS

### Patient population

We examined a patient cohort representative of Boston’s inner-city population comprising 46% African American population. About 60% of the patients were male, about 84% had hypertension, about 75% cardiovascular disease and about 43% had diabetes. About 82% of patients had chronic kidney disease (CKD) stage 3 to 5; 6% had stage 2 CKD, and the rest had stage 1 CKD. About 35% of patients had nephrotic-range proteinuria (>3.5 g/day). On the basis of varied genetic background and several co-morbidities (described above), it is worth noting that the dataset that we generated provides a wide range of glomerular morphologies including cases containing normal glomeruli as well as the ones that manifest partial or global sclerosis.

### Glomerular classification model

The sliding window operator with a small stride (20 pixels) generated a large number of cropped images, and histogram-based thresholding selected many of the images containing the kidney tissue from the ones that contained only the background (Figure S1). This thresholding method was time-efficient and it was able to filter more than half of the cropped images. Using the selected data, binary classification models, constructed by fine-tuning a well-known pre-trained CNN architecture (Inception V3) [10], identified images with a NPS or GS glomerulus with high accuracy across 4 different train-test splits (Table 2). Combining random whitening and data augmentation strategies resulted in better CNN model performance on the testing data as exemplified by model accuracy and kappa (Table 2). The CNN model accuracy on the test data across the four train-test splits ranged from 89.66% to 95.06%. Also, kappa for these cases ranged from 0.8079 to 0.9111.

**Table 2.**
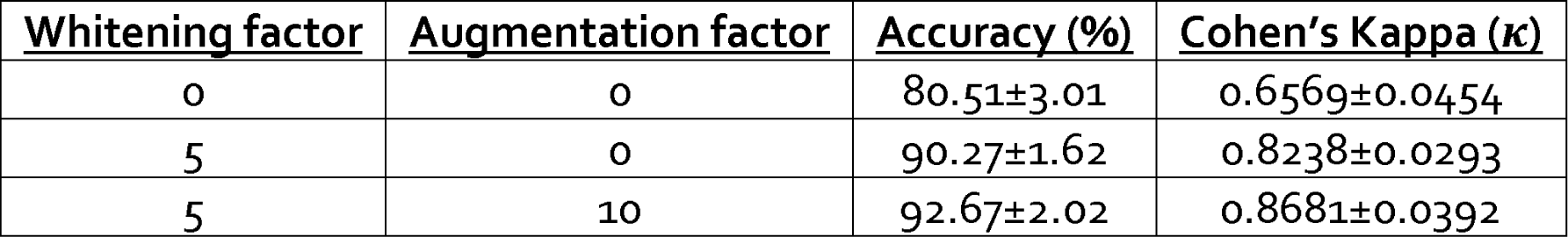
CNN model performance. Three different models were developed to understand the effect of random whitening as well as other data augmentation strategies on the CNN model performance. Model performance is shown on test data that was not used for model training.

| **<u>Whitening factor</u>** | **<u>Augmentation factor</u>** | **<u>Accuracy (%)</u>** | **<u>Cohen's Kappa ($\kappa$)</u>** |
| --- | --- | --- | --- |
| 0 | 0 | 80.51±3.01 | 0.6569±0.0454 |
| 5 | 0 | 90.27±1.62 | 0.8238±0.0293 |
| 5 | 10 | 92.67±2.02 | 0.8681±0.0392 |

### Glomerular segmentation model

The glomerular classification model that generated the best classification performance (Accuracy = 95.06%), was used for segmenting the glomeruli on the test images. The image processing routine involving image binarization, distance transform and the watershed method to segment the GS glomeruli performed well on the test data. Specifically, when the trained CNN model was processed in the form of a sliding window operation on the original images, the model scanned through each region of the image and detected several 300×300×3 pixel windows as non-glomerular regions (Model specificity = 0.999). The image processing routine was also able to identify and mark the GS glomeruli on different test images (Model sensitivity = 0.558, F_1_-score = 0.623, Matthew correlation coefficient = 0.628).

## DISCUSSION

Deep learning algorithms are transforming medicine especially the way by which images and other forms of data are analyzed to uncover interesting patterns and facilitate clinical diagnosis and management of patients [13]. This is especially the case in the field of digital pathology where we and others are employing these powerful techniques to address specific questions in a spectrum of disease scenarios [9, 14–18]. For many of these cases, the clinical workflow is quite similar, i.e., a biopsy procedure is performed to extract a tiny portion of the organ, which is then subjected to a series of histological staining processes prior to the evaluation by a pathologist. In facilities that are equipped with biopsy slide scanners, the tissue slides are digitized to generate images, which then serve as the input data of interest. The deep learning algorithms can read and process these digital signatures to extract relevant quantitative information or associate them with corresponding outputs of interest. Once trained on sufficient number of cases, these models can have the ability to predict on new test cases that the model has never seen before with remarkable accuracy. Currently, the process of biopsy digitization is not performed at all the centers and thus not integrated within all clinical practices as of today. However, there is a growing interest in this direction as the medical community at large is realizing the enormous potential of such a resource. In parallel, efforts should be employed on analyzing these images with cutting-edge techniques to extract maximum benefit for the management of patients. Eventually, this approach will complement the pathologists to improve their accuracy and workflow.

Assessment of renal pathology slides has several features worthy of consideration. The biopsy report methodically deals with all the components of the slide with different staining and clinical correlation is pursued to eventually arrive at a diagnosis. Some of the features (or descriptors) are objective (number of glomeruli), while others are descriptive (type of sclerosis). While the latter item calls upon the expertise of a pathologist, the former can be easily automated. Also, features such as the location (cellular or compartmental) and distribution (focal or diffuse or segmental or global) of the intra-glomerular damage determine the type of glomerular disease (glomerulonephritis vs glomerulosclerosis) can be automated [19–29]. Availability of digitized images provides an immense resource which has opened up opportunities to leverage different tools to improve the analysis of such descriptors that characterize the disease.

Our CNN model identified the presence of an NPS or a GS glomerulus with high accuracy even when a small number of cropped images were used for training (n=1496) (Table 2). It was possible to overcome this limitation with the transfer learning approach. Also, the process of introducing additional noise/variability using random whitening and other data augmentation strategies has shown to limit model overfitting and increase model generalizability (Table 2). Our image analysis pipeline processed the heatmaps to complete the segmentation process that resulted in output images with highlighted areas of the selected GS glomeruli. Note that while simple binarization involves thresholding an image based on a pre-selected value, Otsu’s binarization assumes that an image contains two classes of pixels, and then searches for a threshold that minimizes intra-class variance between the classes [11]. The watershed transformation treats the image it operates upon like a topographic map, with the brightness of each point representing its height, and finds the lines that run along the tops of ridges [12]. Ultimately, the nature of the locations of the glomeruli dictated the performance of the image processing pipeline.

While the current body of work differentiates itself from previous studies [30–33], it is also complementary to them. Integration of these type of studies is needed to generate a comprehensive platform of analyzing digitized renal biopsies using ML techniques. For example, Bukowy et. al. used an ML technique on about 28,000 mouse glomeruli using the periodic acid-Schiff (PAS) stain [33]. While the PAS stain is used for the assessment of basement membrane, the trichrome stain is better suited to evaluate fibrosis and sclerosis. While the use of mouse kidney biopsies facilitate analysis on a large number of glomeruli, it is imperative to examine and validate ML-based findings on human glomeruli for meaningful translation. While the latter is ideal, there may be limitation on the number of samples. Methods such as transfer learning could be used to analyze such small datasets without compromising on the accuracy of the model. Lastly, the cohort used for the current work is ethnically and racially diverse, which adds to the translational significance of these findings.

Our study has the following limitations. We relied on manual observation of cropped images to classify them as the ones containing NPS or GS glomeruli or non-glomerular aspects of the biopsy. A “gold standard” for glomerular identification can help minimize the bias associated with manual selection. Also, our model was only able to discriminate GS glomeruli from normal or partially sclerosed ones. While discriminating normal from partially sclerosed ones also can be helpful, a routinely assessed pathologic finding is the percent glomerulosclerosis, and we believe our current model can successfully estimate this value on a digitized kidney biopsy image. In future, we plan to develop a model that can distinguish between normal and partially sclerosed glomeruli. This task can be accomplished successfully when we have access to a large collection of NPS and GS glomerular images.

Our paradigm for identifying and segmenting glomeruli is likely to be useful for the processing of the pathology slides and can be easily extended to images generated using other staining protocols. Adoption of such methods without disturbing the workflow of a pathologist can expedite the assessment of slides and serve as a first step toward more comprehensive, automated analysis. Further validation of the deep learning framework along with the image processing operations across different clinical practices and image datasets is necessary to validate this technique across the full distribution and spectrum of lesions encountered in a typical nephropathology service.

## ACKNOWLEDGMENTS

This project was supported in part by the National Center for Advancing Translational Sciences, National Institutes of Health, through BU-CTSI Grant (1UL1TR001430), a Scientist Development Grant (17SDG33670323) from the American Heart Association and Boston University’s Digital Health Initiative Research Incubation Award (#2018-02-008) to VBK, Boston University’s Undergraduate Research Opportunities Program funding to LAM, and NIH grants (R01-HL132325 and R01-CA175382) to VCC.

## SUPPLEMENTARY MATERIAL

**Supplement:** Glomerular dataset generation, Model training, Data augmentation

**Figure S1.**
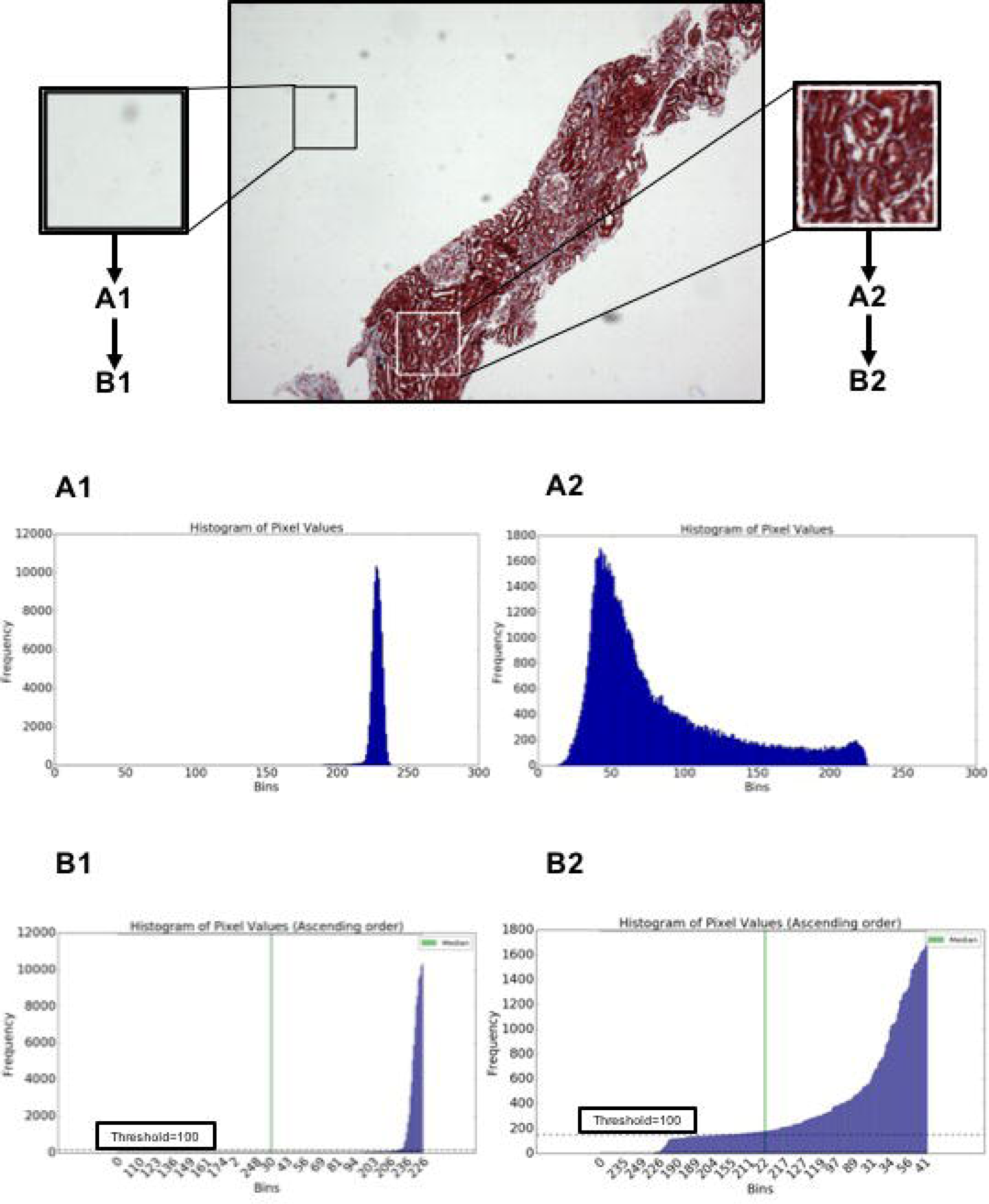
Histogram-based thresholding. A sliding window operator scanned the entire original image of size 2560×1920×3 pixels and generated cropped images of size 300×300×3 pixels. For each cropped image, a histogram based on pixel intensity was generated (**A_1_** for a cropped image representing the background and **A_2_** representing a portion of the kidney biopsy). These histograms were then reordered according to the bin frequency. A threshold value of 100 was empirically selected as a cutoff and median value for the bin frequency was computed. Images with a median value below the cutoff were selected as the ones representing the background (**B_1_**), and the ones with a median value above the cutoff were selected as part of the kidney biopsy (**B_2_**).

