## Supplement for "Segmentation of Glomeruli Within Trichrome Images Using Deep Learning"

***Glomerular dataset generation***

The size of the cropped images was empirically chosen such that any glomerulus observed within the original 40x images was able to fit well within the bounds of the cropped image size. For each original image, the sliding window operator began at the top left corner of the 40x image and moved right with a stride of 20 pixels after cropping the 300x300x3 window. The stride size was empirically determined such that it resulted in each glomerulus being captured completely in at least one cropped image. In total, this process generated about over 100,000 unique images of size 300x300x3 from 275 unique 40x images of size 2560x1920x3 pixels. In order to expedite the cropping process, windows consisting of purely non-biopsy portions (i.e. background) were automatically ignored. For each cropped segment, we computed the median intensity of all the pixels and selected only those cases with a median bin frequency value lower than an empirically estimated threshold intensity of 100 (**Figure S1**). Our idea of thresholding was based on selecting a single cutoff value for the bin frequency (=100). The underlying assumption was that an image with only the background had a higher frequency of pixels with the same intensity values (**A1** in **Figure S1**), whereas a cropped image with tissue within it had a more distributed range of intensity values with lower frequencies (**A2** in **Figure S1**). In order to compute the median, the histogram bins (x-axis) were arranged in the ascending order of their corresponding frequencies (y-axis). In case of the background segment, since the frequencies were concentrated in a very narrow bin range when arranged in the ascending order, the median turned out to be lesser than the threshold value of frequency (=100) (**B1** in **Figure S1**). For the tissue segments, since the frequencies are distributed over a range of bins, when arranged in ascending order, the median frequency was found to be higher than the threshold value of frequency (=100) (**B2** in **Figure S1**). Following histogram-based thresholding, we manually selected a unique set of glomerular and non-glomerular images from them that were then used for CNN model development. This process selected about 40,000 images from over 100,000 images. We then examined all the non-background images and then selected images that included a unique glomerulus (n=745). An equal number of images were selected from the remaining non-background images with non-glomerular tissue as control cases. Together, these images formed the training data, with an output label of ‘0’ assigned to the non-glomerular images, ‘1’ for normal or partially sclerosed glomerular images and ‘2’ for globally sclerosed glomerular images. Note that the non-glomerular images were selected across different portions of the biopsy in order to capture variability within a patient and to include several unique aspects of the biopsy (such as tubular elements, interstitial spaces, vascular regions, etc.).

***Model training***

Our CNN model was trained using back-propagation. Using the framework of transfer learning, we first trained only the top layers that were randomly initialized by freezing all the convolutional layers. We then trained the model using our data for several epochs with the early stopping criteria that monitored the validation loss. We used “rmsprop” with a decay of 0.9, a momentum of 0.9, an epsilon of 0.1, along with the use of L_1_ and L_2_ hybrid regularizers (0.01, 0.01). After the top layers were trained, we performed fine-tuning of the convolutional layers by freezing the bottom (N) layers and training the remaining top layers. For this task, we used stochastic gradient descent with a learning rate of 0.001, a momentum of 0.9, and L_1_ and L_2_ regularizers (0.01, 0.01). We finally trained the model again by fine-tuning the top blocks alongside the top dense layers. We used Google’s TensorFlow (<https://www.tensorflow.org>) back-end to train, validate, and test our network.

***Data augmentation***

During training, images were augmented by a factor of 10, therefore creating 10 additional copies per image. Each copy of the image was then rotated randomly between 0° and 180°, randomly shifted in horizontal and vertical directions, shifted the color channels, sheared, and zoomed, all by a scale of 0.1. Images were also flipped vertically and horizontally, with a probability of 0.5. The expectation was that data augmentation in this fashion was able to minimize model ‘overfitting’, and thus enhanced the generalizability of these models.
